# The synergic role of actomyosin architecture and biased detachment in muscle energetics: insights in cross bridge mechanism beyond the lever-arm swing

**DOI:** 10.1101/2021.03.23.436693

**Authors:** Lorenzo Marcucci, Hiroki Fukunaga, Toshio Yanagida, Mitsuhiro Iwaki

**Author notes:** **Corresponding Author:** Lorenzo Marcucci, **Email**. **Author Contributions:** L.M. and M.I. designed the project; L.M. developed the theoretical formalism, defined the mathematical model and performed the numerical simulations and numerical data analysis; M.I. established the experimental protocols; H.F. and M.I. contributed new reagents/analytic tools; H.F. analysed the experimental data; T.Y. supervised the project; L.M. wrote the paper with input from M.I. **Competing Interest Statement:** The authors declare no competing interests.

## Abstract

Muscle energetics reflects the ability of myosin motors to convert chemical energy into mechanical energy. How this process takes place remains one of the most elusive questions in the field. Here we combined experimental measurements of in vitro sliding velocity based on DNA-origami built filaments carrying myosins with different lever arm length and simulations based on a Monte-Carlo model which accounts for three basic components: (i) the geometrical hindrance, (ii) the mechano-sensing mechanism, and (iii) the biased kinetics for stretched or compressed motors. The model simulations showed that the geometrical hindrance due to acto-myosin spatial mismatching and the preferential detachment of compressed motors are synergic in generating the rapid increase in the ATP-ase rate from isometric to moderate velocities of contraction, thus acting as an energy-conservation strategy in muscle contraction. The velocity measurements on a DNA-origami filament that preserves the motors’ distribution showed that geometrical hindrance and biased detachment generate a non-zero sliding velocity even without rotation of the myosin lever-arm, which is widely recognized as the basic event in muscle contraction. Because biased detachment is a mechanism for the rectification of thermal fluctuations, in the Brownian-ratchet framework, we predict that it requires a non-negligible amount of energy to preserve the second law of thermodynamics. Taken together, our theoretical and experimental results elucidate non-conventional components in the chemo-mechanical energy transduction in muscle.

## Introduction

The smallest contractile unit in muscle, the half-sarcomere, reduces its length through the sliding of thin filaments, which are formed by actin monomers, relatively to thick filaments, which are formed by a structured arrangement of myosin II motors [1–3]. It is widely accepted that this relative motion is generated by the cyclical interaction of myosin motors with actin monomers, which includes attachment, biased rotation of the lever-arm, which generates force, and detachment (Figure 1A, “active cycle”) [4]. This process is powered by hydrolyzation of the high energy molecule adenosine triphosphate (ATP) into adenosine diphosphate (ADP) and phosphate (Pi). Macroscopic power is the result of myosin motors converting ATP chemical energy into mechanical energy. How this process takes place is one of the most elusive questions in this field.

**Figure 1.**
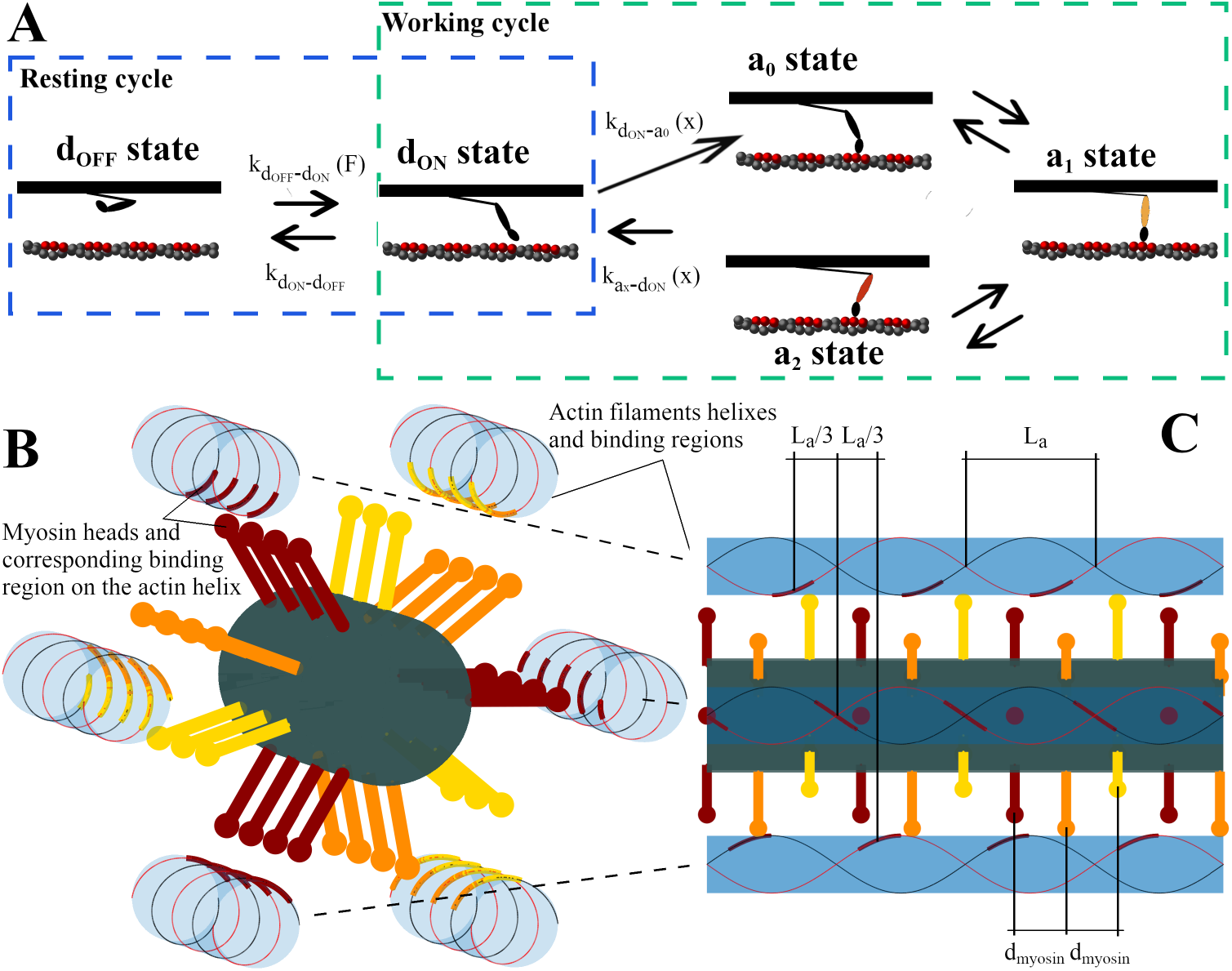
Cross-bridge cycle and pseudo-replication of the 3D structure in the model. (A): The cross- bridge cycle where, in an otherwise classical model, the super-relaxed state in myosin motors is taken into account, splitting the whole cycle into a resting cycle for the activation of the thick filament and a working cycle, where a two-step power stroke generates force. The activation rate depends on the force acting on the thick filament. The attachment and detachment rates depend on the relative position of the motor with respect to its anchor. (B): 3D structure of one thick filament surrounded by six thin filaments. 4 crowns of motors are shown. The two helixes on the thin filaments are indicated along with the available binding regions, highlighted with the color of the corresponding motor heads. C) Lateral view of the 3D structure. Only the binding regions for the deep red motors are indicated, showing *L*_*a*_/3 shifts. The mismatch between actin-binding regions and the motors can be particularly appreciated in the motors pointing out of the figure plane toward the viewer (deep red dots in the middle of the figure) and the corresponding binding regions. The first head from the left is pointing exactly to the center of the binding region. The second is more displaced but still within it. The third and fourth heads cannot attach because of the geometrical hindrance.

The muscle power output *P* = *Fν* generated by a muscle fiber during isotonic contraction, i.e. at a steady velocity *ν* against a submaximal external force *F*, reaches its maximum *P*_*max*_ at about 1/3 of the maximum velocity of contraction *ν*_*max*_ [5]. The relatively high *P*_*max*_ must be associated to a relatively high rate of attachment by the myosin motors. In this way, fresh motors can sustain the high force as others become exhausted by the sliding of the filaments. However, when introduced in quantitative mathematical models, this high attachment rate is in apparent contrast with at least three other features related to muscle energetics. As a first discrepancy, it would produce a faster rise in force during activation than that observed experimentally. This limit was highlighted in the first mathematical model of the *F*(*ν*) curve proposed by A.F. Huxley in 1957 [6]. Moreover, a high attachment rate is optimized for intermediate forces but is a waste of energy for isometric contraction against maximum force *F*_*max*_ or during an unloaded contraction at maximum velocity *ν*_*max*_, because it would generate a high ATP consumption rate (*r*_*ATP*_). Instead, as a second discrepancy, the ATP consumption rate in isometric contraction is kept as low as 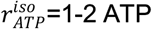 per myosin motor per second, and consequently the *r*_*ATP*_(*ν*) relationship has a steep behavior, increasing by a factor of about 5 to its maximum value 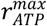 [5,7–9]. Notably, the modulation of the detachment rate in isometric contraction is limited by experimental evidence that showed about 30% of myosin motors are attached to actin [9] in this condition. Finally, despite technical difficulties to precisely estimate *r*_*ATP*_(*ν*) at higher velocities of contraction, its first derivative clearly decreases, and possibly *r*_*ATP*_ does too in absolute values [10] as *ν* approaches *ν*_*max*_, resulting in the third discrepancy. In this manuscript, we investigated these energy-conserving strategies through *in silico* (model simulation) and *in vitro* (motility assay with DNA origami filaments) methods.

Different hypotheses have been proposed to solve one or more of the discrepancies with mathematical models (see below in this section, [11] and references therein). A recent model, proposed by one of the authors [11] supported a possible role for the activation of the thick filament through the mechano-sensing mechanism [12], described as follows. Detached motors exist in at least two stable states in relaxed muscles [13]: a detached, super-relaxed, or OFF, state (*d*_*OFF*_), and a detached but active, or ON, state (*d*_*ON*_). In the OFF state, myosin S1 sub-fragments lie down along the thick backbone, while in the ON state myosin motors point to the actin filament ready to start the active cycle (Figure 1A). Experimental studies suggest that the contraction force is modulated not only by classic [Ca^2+^]-mediated activation of the thin filament, but also by force- mediated activation of the thick filament [12,14,15]. In fact, the amount of ON motors is proportional to the force sustained by the filament itself, and this effect has important roles in muscle contraction [16–20].

Introducing the mechano-sensing mechanism into an otherwise classic model [11], it is possible to preserve *P*_*max*_ and to solve the first and third discrepancies, but not to explain the high 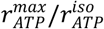 ratio. In particular, the model is not able to keep 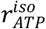 low, because at tetanic isometric conditions all the myosin motors are ON and ready to consume ATP.

To modulate the ATP consumption at different velocities of contraction, something must modify the real or apparent rates of formation and dissociation of actomyosin complex, the cyclical interaction which consumes ATP. In the model these are described by the attachment and detachment rates between the detached active state *d*_*ON*_ and each of the attached states *a*_*x*_ (Figure 1A). At least two experimentally verified components influence the apparent attachment and detachment rates at different *ν*: the preferential detachment of compressed myosin motors [21,22] and the existence of a limited binding region on the double-helix conformation of the actin filament, as suggested by single molecule experiments [23] and more recently by fast AFM images [24]. The latter component generates geometrical hindrance to the actomyosin interaction due to the spatial mismatch between myosin motors and actin filament periodicities (Figure 1B and C). Both components influence the amount of motors ready to work during the relative sliding.

Some models have successfully simulated the high 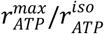 ratio, including the original model proposed by A.F. Huxley in 1957 [6]. However, despite this model hypothesizing for the first time and without any experimental insights the preferential detachment in negatively strained motors, it is not present in the definition of the total rate of energy liberation (eq. 5 in the original paper) used to fit the experimental *r*_*ATP*_(*ν*) from [25]. This is a mathematical consequence of the imposed condition that at each strain of the motors, *x*, the sum of state probabilities is equal to one. This condition doesn’t account for the preservation of the finite number of motors and target regions on the actin filament. This problem is common in mean-field models, which average the behavior of populations of myosin motors, and comes from the fact that they focus only on the binding region and therefore don’t account for the motors that move out of this region. Piazzesi and Lombardi introduced a periodic condition at the boundary of the attachment region in order to preserve the number of myosin motors (but not the actin target regions availability). They obtained a high 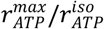 ratio, hypothesizing a second “short” path in the cross-bridge cycle [8], which is characterized by faster attachment-detachment rates. A high 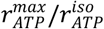 ratio can also be obtained by hypothesizing that attached myosin motors can slip on the actin filament across two consecutive actin monomers [26] or by imposing a dependence of the attachment rate on the velocity of contraction [27]. However, these hypotheses have not been verified experimentally. Moreover, in the mean-field models, the geometrical hindrance cannot be easily introduced within the constraints to preserve the total number of myosin motors. The Monte-Carlo approach is more suitable to study the effects of the geometry, since it analyzes the behavior of each motor independently. However, few models based on this technique include an analysis of 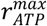 or analyze other above-mentioned incongruences. For instance, Smith and Mijailovich obtained a good fitting of *r*_*ATP*_(*ν*) but by accepting a fast rate of force generation during activation (7 ms of the half-rise time) [28].

Despite both the preferential detachment and the geometrical hindrance being recognized to influence the *r*_*ATP*_(*ν*) relationship, to the best of our knowledge, no previous work has assessed quantitatively their relevance. In particular, assessing the role of the preferential detachment is of importance in muscle energetics, because, as a Brownian ratchet mechanism which rectifies thermal fluctuations, it requires ATP energy to preserve the second law of thermodynamics. This amount of energy would not be available for the lever-arm rotation. Our previous model [11] accounted for the preferential detachment but not for the geometrical hindrance and showed a mild effect on the *r*_*ATP*_(*ν*) relationship from zero to moderate velocities of contraction. Experimentally, Spudich and coworkers showed that the sliding velocity of an actin filament on a bed of myosin motors linearly depends on the lever-arm length and inferred a zero velocity in the absence of any lever-arm rotation [29]. These results may suggest that the energy associated with the preferential detachment is negligible. However, we show here that this conclusion cannot be valid when the muscle structure is preserved.

In this study, we used a Monte-Carlo model which accounts for three basic components: (i) the geometrical hindrance, (ii) the mechano-sensing mechanism, and (iii) the biased kinetics for stretched or compressed motors. These components are derived from experimental observations, limiting the use of unverified hypotheses. We showed that the geometrical hindrance and the biased detachment synergistically act to reproduce the steep increase of the ATP-ase rate. We were able to confirm the model prediction with in vitro motility experiments, by extrapolating the relative sliding velocity of two filaments at different lever-arm lengths, with the spatial mismatch between the actin and myosin filament periodicities preserved. Our data showed that a non-negligible velocity is present even in the absence of the lever-arm, suggesting a non-conventional component in the chemo-mechanical energy transduction in muscle.

## Results

### Geometrical hindrance limits ATP consumption in isometric conditions and at low but not intermediate velocities of contraction

We first test the effect of geometrical hindrance (basic component (i)) by comparing the predicted *r*_*ATP*_(*ν*) in models where the periodic target regions on the actin filament are progressively reduced, starting from a situation where myosin heads can attach everywhere along the thin filament. All other hypotheses are the same, including the mechano-sensing mechanism, and the biased kinetics for stretched or compressed motors (basic components (ii) and (iii)). To produce a coherent comparison, we modulate the parameters of each model to fit three experimentally observed data: the *F*(*ν*) curve (Figure 2A, experimental data from *Rana temporaria* [9]), the kinetics of force generation in activation and after a small shortening (Figure 2B, experimental data from *rana temporaria* [12]), and the amount of attached heads in isometric activation (about 30%, experimental data from *Rana temporaria* [9], see SI Figure S1). Notably, our numerical simulations predict that a minimum of three actin monomers should be available for actomyosin interactions in order to fulfil the last request (SI Figure S1). Therefore, in the rest of this manuscript, we used the 3-actins model to infer the influence of the geometrical organization on the ATP consumption.

**Figure 2.**
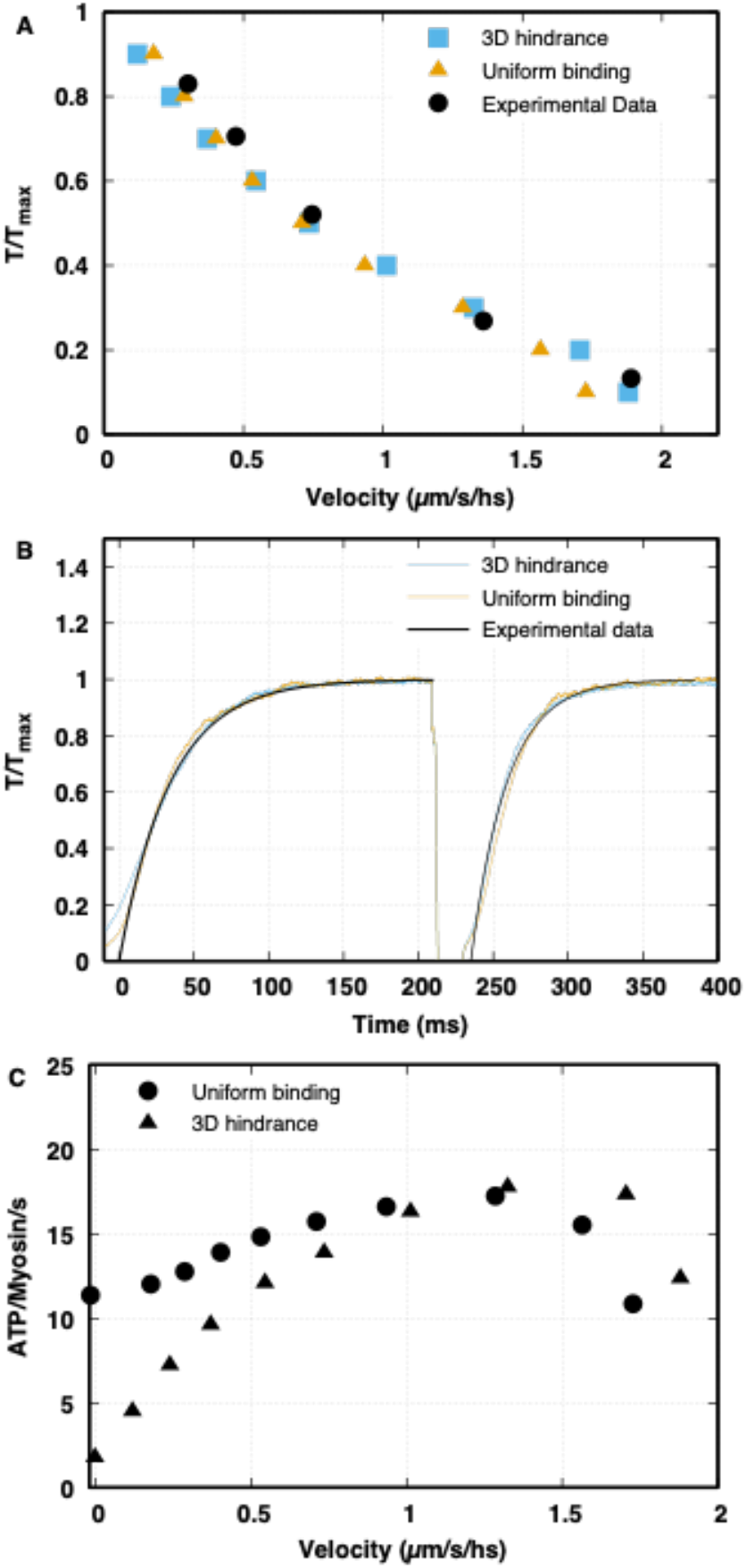
Comparison of the uniform binding region model vs. the limited target zones model, which generates geometrical hindrance. (A) The attachment rate 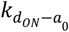 is adjusted to fit the *F*(*ν*) curve (experimental data for *Rana temporaria* from [9], velocity in µm per second per half sarcomere (hs)). Then, the detachment rate 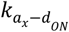 is adjusted to have 30% of motors attached in isometric contraction (experimental data for *Rana temporaria* from [9]). (B) The rate of activation of the thick filament as a function of the force 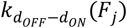 is adjusted to fit the kinetics of the force generation during activation and force recovery after an unloaded shortening (experimental data for *Rana temporaria* from [12]). (C) Despite the mechanical behaviors being very similar for the two models, the uniformly distributed target zones (circles) induce a high 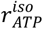, while the limited target zones drastically reduce it (triangles).

Despite us imposing similar mechanical behaviors, the predicted 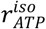 is greatly affected by the extension of the target zones along the thin filament (Figure 2C). In the case of continuous suitable binding sites along the thin filaments, the predicted 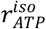 is about 12 ATP molecules per myosin head per second, much higher than experimental estimations, leading to a maximum-to-isometric ratio, 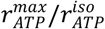, of 1.5 (Figure 2C circles). It is important to note here that this low 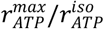 ratio is obtained in the presence of a preferential detachment of compressed motors.

When a reduced range of target zones is imposed, the emerging geometrical hindrance limits the attachment of the myosin motors in isometric contraction, and 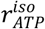 drops to more physiological values (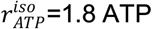 per myosin motor per second, Figure 2C triangles).

However, when the applied external force is lower than *F*_*max*_, the contraction occurs, and the relative sliding of the filaments makes new target zones available to the detached heads, increasing the ATP-ase rate. At intermediate velocities of contraction, the effect of the geometrical hindrance almost disappears, leading to a predicted 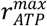 similar to that in the continuous model. In the end, the reduced target zones model shows a sharp increase of *r*_*ATP*_(*ν*), closer to that observed experimentally. The 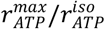 ratio in this case reaches a value of 9, which is even higher than the experimental data. However, in our analysis, the model is kept relatively simple to focus on the three basic components (see Discussion section). Therefore, we show that preferential detachment alone is not sufficient to explain the 5-fold increase in *r*_*ATP*_ from isometric to maximum values, with this increase limited to 50% in our simulations. Instead, we show that the preferential detachment and the geometrical hindrance have the potential to fully explain the observed increase.

Looking at the *r*_*ATP*_(*ν*) relationship at higher velocities, both models predict a drop in the ATP-ase rate due to the mechano-sensing mechanism. Sensing a low force along the thick filament, when the external force is reduced, several myosin motors switch off, exiting from the “active” ATP-consuming cycle. The mechano-sensing mechanism here becomes a second energy-conserving sensor. This result is consistent with the observation that the number of myosins needed to drive an unloaded contraction at maximum velocity is small [30].

### Preferential detachment has a synergistic effect with geometrical hindrance to increase the dependence of ATP-ase activity on the velocity of contraction

The models used in the previous section are both based on Huxley’s original 1957 hypotheses for preferential detachment of the myosin motors when compressed and a preferential attachment when stretched [6]. Biased kinetics without geometrical hindrance cannot reproduce a high 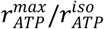 ratio (Figure 2, circles). On the other hand, we test here if the geometrical hindrance without biased kinetics can fully account for it.

We approach this question by first excluding from the model both geometrical hindrance and biased kinetics (basic components (i) and (iii)) to verify that no other dependence of *r*_*ATP*_ on *ν* is present (Figure 3, upper panel, squares). As expected, this case predicts a constant *r*_*ATP*_ from isometric conditions to intermediate velocities of contraction, still showing the decrease at low external forces due to the mechano-sensing mechanism. Reintroducing only the discontinuous target zones on the actin filament, an increase in *r*_*ATP*_(*ν*) appears again at low *ν*, but the 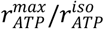 ratio is only 1.2, much lower than what was observed experimentally (Figure 3, upper panel, circles). Whilst some adjustment in the parameters are required to make the analysis comparable (see SI text), this result clearly shows that the geometrical hindrance component alone cannot explain the high values of the 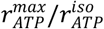 ratio, which in this model can increase only about 20%.

**Figure 3.**
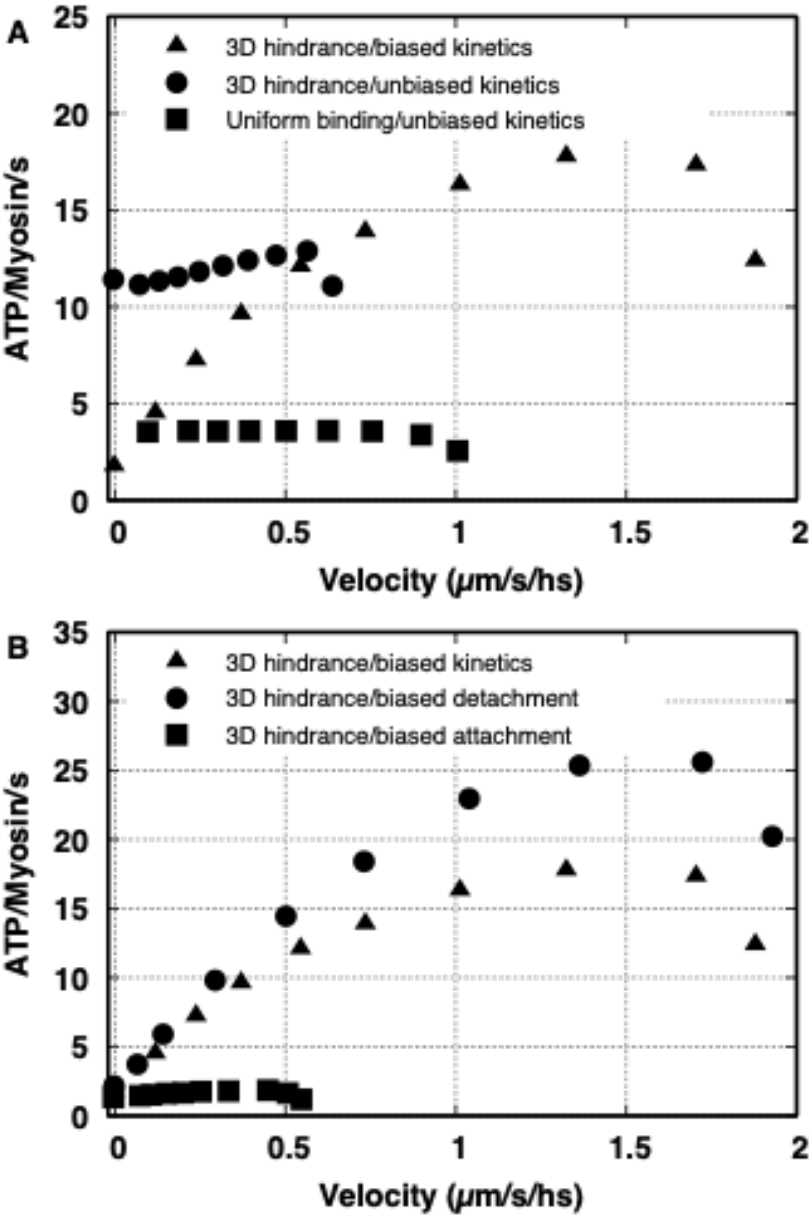
Analysis of the effect of biased kinetics and geometrical hindrance on *r*_*ATP*_(*ν*). Triangles refer to the model with limited target zones and biased kinetics for comparison. (A) Both the limited target zones and the preferential detachment are excluded (squares): the ATP-ase rate doesn’t depend on the velocity of contraction except at low external forces (high velocity) due to the mechano-sensing mechanism. When the limited target zone is present without the effect of the preferential detachment (circles), the r_ATP_^max^/r_ATP_^iso^ ratio is lower than the experimental value. (B) The geometrical hindrance is included without biased attachment (circles) or without biased detachment (squares). The high r_ATP_^max^/r_ATP_^iso^ ratio is maintained only when the geometrical hindrance acts together with the preferential detachment of myosin motors. On the contrary, the preferential attachment has no effect on the *r*_*ATP*_(*ν*) relationship.

Next, we explore how much preferential detachment and attachment separately influence the 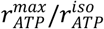 ratio when the geometrical hindrance is included. Both properties have been introduced theoretically in [6] and, more recently, gained experimental support [21,22,24,31]. We first note that eliminating the preferential attachment (Figure 3, lower panel, circles) does not affect significantly the 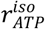 ratio, but 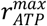 maintains values even higher than in the previous analysis. This is due to the different parameters needed to match the *F*(*ν*) curve in the absence of the preferential attachment (see SI Figure S2). On the contrary, excluding only the preferential detachment, the 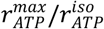 ratio drops to low values (Figure 3, lower panel, squares, and see SI text). Also, *r*_*ATP*_ is constant with *ν* even in the presence of the preferential attachment without the geometrical hindrance (not shown), which confirms that it doesn’t introduce any dependence for *r*_*ATP*_ on the relative position of the two filaments and thus on the velocity.

To summarize, as one may expect, the preferential detachment and the geometrical hindrance are crucial in the 500% increase of 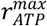 with respect to 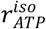, but our simulations suggest also that the two effects, which individually account for an increase of 50% and 20%, respectively, must act synergistically to fully explain the experimental data.

### Thermal ratchet components can account for up to 15% of maximum velocity in physiological conditions

Biased kinetics break the detailed balance [32–34], and to preserve the second law of thermodynamics, they require an external source of energy, namely ATP energy. This energy is needed to rectify thermal fluctuations, making muscles partially act as a Brownian ratchet, and are not available to prompt rotation of the lever-arm. However, this role may be appreciated only when considered in combination with the geometrical hindrance, as suggested from our simulations.

We tested the model predictions directly by observing the *in vitro* effect of biased kinetics when the acto-myosin spacing is preserved. To achieve our goal, we used the DNA-origami technique [24,35], a nanotechnology suitable to construct arbitrary three-dimensional nanostructures from DNA. Ten DNA duplexes were bundled in parallel [24] to build a rod nanostructure (DNA rod) to which we attached myosin motors with precise spacing of 14.3 nm (Figure 4A). The monomeric DNA rod (∼250 nm in length) can attach 18 myosin motors with the same orientation, and the rod was oligomerized to form an ∼ 1 µm-long filament structure (tetramer or pentamer). To observe actin sliding motion along our DNA rod-myosin complex, we first attached the complex onto a coverslip and then introduced actin filaments with ATP. We simultaneously observed the fluorescently labelled (TAMRA) DNA rod and actin filament (ATTO647N) (Supplementary movie 1) and confirmed the actin filaments sliding over a DNA rod are longer than the DNA rod itself, suggesting that the number of interacting myosin heads was constant. Then, to precisely determine the actin sliding velocity, we attached fluorescent quantum dots (QDs) to the filaments and tracked the fluorescent spots with a few nanometer accuracy [36] using total internal reflection fluorescence microscopy (TIRFM). The actin filament constantly moved in one direction, and the velocity depended on the myosin constructs (Figure 4B).

**Figure 4.**
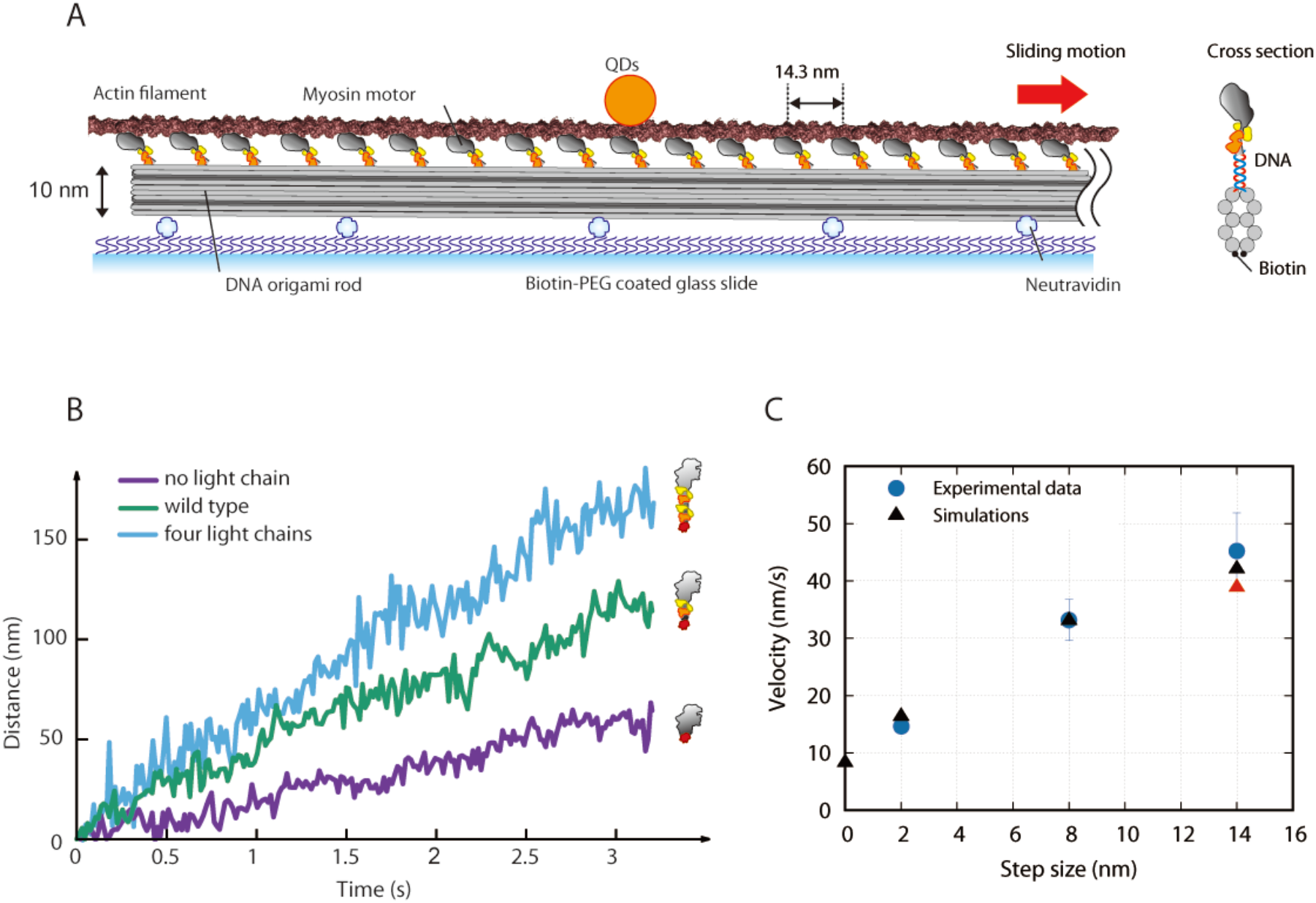
*In vitro* actin sliding assay driven by a DNA rod-myosin complex. (A) Schematic of the in vitro sliding assay. Qdots (QDs) were attached to an actin filament, and a DNA rod-myosin complex was adsorbed onto a glass slide. Biotins were attached to the bottom of the DNA rod to fix the rod to the neutravidin-coated glass slide, and DNA handles (21-base single-stranded DNA) were attached to the top of the DNA rod (see cross section shown in right cartoon) at 14.3 nm spacing along the DNA rod. Because anti-handles were labeled to myosins, the myosins were precisely positioned to the DNA rods with a spacing of 14.3 nm. ATP concentration, 0.5 µM. (B) Trajectories of the actin sliding distance for three distinct myosin mutants. Purple, no light chains (without lever-arm); Green, WT; Blue, four light chains (double-length lever-arm). (C) Predicted average velocities of the sliding model (black triangles) at different step sizes and the corresponding velocities observed experimentally (blue dots) are related to different lengths of the lever-arm. The intercept at zero step-size is predicted by the model preventing the motor to swing the lever-arm. Experimental data was obtained from at least 12 independent experiments and n=83, 77 and 72 filaments for no light chain, wild type and four light chains myosin, respectively. Error bars indicate standard deviation.

We designed myosin mutants with three different lever-arm lengths: no light chains, wild type (WT) and four light chains (see Material and Methods and SI Figure S3). Our data showed that the velocity increases with increasing lever-arm length (Figure 4C blue dots): 15.5±5.6 (mean±s.d.) nm/s for no light chains, 35.0±8.6 nm/s for WT, and 42.7±17.7 nm/s for four light chains myosin.

Even in the mutant without light chains, a non-zero velocity was previously observed experimentally in the absence of preserved spatial mismatching and associated to the putative fulcrum of the lever-arm outside the light chains region [29]. However, in our experimental analysis, the relative drop in the sliding speed of a system without light chains with respect to WT is about 56%, which is much less pronounced than in previous experimental data based on a bed of myosin motors randomly distributed (drop of about 80% [29]). Consequently, in our setup, the estimated velocity at the intercept with no lever-arm rotation is not negligible. To support the experimentally inferred velocity in this extreme case, we use an *in silico* approach and simulate the velocity in a system where the lever-arm rotation is completely prevented (see Material and Methods and SI text). To maximize the predictive power of this model, we use the detachment rate experimentally observed in [21] through Bell’s equation (see Material and Methods). Again, the parameters are adjusted to match the force generation and regeneration, the force velocity curve, the percentage of attached heads in isometric contraction, and *r*_*ATP*_(*ν*) (SI figure S4). The constant parameter that simulates the effect of the ATP concentration (see Materials and Methods) was used to fit the sliding velocity for the WT case.

The simulated sliding velocities for different power stroke sizes (all the way to zero) are reported in Figure 4C (triangles) and superimposed to the corresponding experimental data. When the lever-arm is prevented to rotate from the pre-power stroke state, the model predicts a sliding velocity of 8.6 nm/s, which is about 25% the sliding velocity for the WT case. Using the same parameters in the whole fiber model, the maximum velocity preventing the lever arm rotation is 0.38 µm/s/hs, which corresponds to 16% of *ν*_*max*_ in the WT case (2.31 µm/s/hs).

In conclusion, we have shown experimentally the ability of motors without a lever-arm swing to preserve velocity in the filament sliding assay, and so, that the energy associated with the preferential detachment is non-negligible, as indicated from our numerical simulation.

## Discussion

Experimental work has shown that muscle, as a chemo-mechanical energy transducer, adapts its energy consumption to external conditions, keeping it low in isometric contraction, but increasing it when a shortening against a submaximal force is required. A car can remain stationary on a slope either by regulating the motor force to be equal to the gravity force, thus consuming energy, or using the brakes, thus saving energy. However, this is a choice made by the driver or, in nano- machines, by a pre-programmed computer. In muscle, this energy-conservation strategy must come naturally from the actomyosin cycle properties. In this manuscript we analyzed the different components responsible for these properties.

Qualitatively, it is quite intuitive how *r*_*ATP*_(*ν*) is regulated by the geometrical hindrance and the preferential detachment. The geometrical hindrance limits some myosin motors to reach the actin target region during isometric contraction, generating a population of detached ON motors that can be used when the two filaments slide past each other. However, this population is rapidly exhausted without a preferential detachment in compressed motors, and in our model the 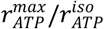 ratio is increased by only 20%. On the other hand, the preferential detachment increases the apparent detachment rate when the relative sliding of the filaments takes place, leading to a higher ATP consumption. However, the increase is proportional to the ratio between the detachment rates in positive and negative stretches, which is constrained by the observed maximum velocity and the fraction of attached motors in isometric conditions. The 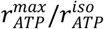 ratio increases by only 50% because of this mechanism in our model. The simulations presented in this manuscript indicate, for the first time, that the 5 folds of the 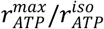 ratio is compatible with other observations of muscle energetics when both the geometrical hindrance and preferential detachment act together, prompting a synergistic effect.

This new insight suggests that the role of the preferential detachment in muscle energetics may be non-negligible only when considered in the physiological muscle architecture. In fact, while the geometrical component doesn’t break the principle of detailed balance, the preferential detachment does. The swinging lever-arm theory describes the contraction of muscle through the movement of the lever arm in a biased direction: myosin motors always attach in the high-energy state of the pre-power stroke and statistically go to the lower-energy state(s) of the post-power stroke(s), requiring ATP energy. This picture limits the importance of the thermal fluctuations, because the process is almost deterministic in its main features. Instead, biased kinetics can be better described in the framework of Brownian ratchets [32–34]. A simple case is represented by the preferential attachment: thermal fluctuations of detached motors occur randomly, but a sensor exists and is able to rectify them, allowing attachment only when the motors are stretched in one direction. This generates a biased force or motion. Similarly, preferential detachment is able to break the detailed balance generating a population of attached motors that are statistically more stretched in one direction, again generating force or motion.

We then experimentally estimate of the unloaded motion associated to the biased kinetics in the absence of the lever-arm rotation while preserving the synergic effect related to the mismatch of the actin and myosin spacing. The effect of the geometrical hindrance on the energetics of muscle contraction cannot be easily analyzed *in vivo* or *in situ* due to the complexity of the system, but it is also elusive in single molecule experiments, because the spatial mismatch is completely lost. Moreover, it is known that myosin filaments in not only the classical *in vitro* actin sliding assay [29] but also a conventional reconstitution technique [37] do not mimic the symmetric bipolar filaments present in sarcomeres and instead assemble randomly [38]. In contrast, our DNA origami-based myosin filaments can control the myosin spacing at less than one nanometer precision to preserve the mismatch. Our setup shows that, when the spatial mismatch is preserved, the drop of the sliding velocity for the mutant without light chains relative to WT was less pronounced than in previous experimental data based on a bed of myosin motors randomly distributed [29]. We speculate that the random distribution of the motors and the loss of the physiological spatial mismatch may hide the effect of the biased actomyosin kinetics. Our results further show that a non-zero velocity is present in the complete absence of the lever-arm rotation, which the model suggests can be attributed to the biased kinetics. Our experimental setup is limited in that it cannot fully estimate the energetics associated to the components not related to the lever-arm rotation, because we cannot estimate the force generated. Therefore, we inferred the relative importance of the biased kinetics on muscle energetics from the 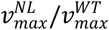 ratio between the maximum velocity of a system with no lever-arm myosin and WT myosin. We found up to 15-25% of the maximum velocity may be attributed to the biased kinetics depending on the [ATP].

Additionally, the mechano-sensing mechanism induces a reduction of active motors during contractions against low forces toward the minimum number needed for the unloaded contraction [30]. Consequently, *r*_*ATP*_(*ν*) shows a decrease toward v_max_. This behavior was observed experimentally by A.V. Hill in 1964 [10]. While other hypotheses are able to reproduce it (see [39] and ref. therein), the mechano-sensing mechanism represents a relatively simple explanation.

We must acknowledge the limits of our approach. Some are shared with any mathematical model and are due to the limited knowledge of crucial parameters in the acto-myosin interaction. Examples are the possible non-linearity of the myosin motors stiffness [40–42], the exact rate of preferential detachment, which was observed in cardiac myosin II motors [21,22] and far from *in vivo* conditions, and the rate of preferential attachment, which has been directly observed for myosin VI [31] and was recently suggested for myosin II [24], but not in a quantitative way. In addition, our model has some intrinsic limits such as inextensible thin and thick filaments and a pseudo-replication of the 3D geometry that may not be accurate. Moreover, we did not include certain debated effects such as myosin cooperativity in the thin filament activation, MyBP-C effects on the stabilization of the OFF state, and calcium-diffusion effects [43–45]. Finally, modifications of the myosin constructs in the experimental setup may affect both the myosin stiffness [46] (see also SI text and Figure S5) and the ability of myosin to interact with actin monomers, though it is unlikely that the modifications would increase the ability. However, despite these limits, the driving hypotheses in the model are mainly based on experimental evidence. Other previous theoretical works showed the importance of accounting for the complex muscle ultra-structure [47–50], while mass action models have attempted to represent the ultra-structure by considering different extensions of the target zones [8,26,51,52]. However, since our model includes only experimentally based hypotheses, it can be used to systematically analyze the dependence of the *r*_*ATP*_(*ν*) curve on the different components, especially in association with the mechano-sensing mechanism, which is a first. We believe that the synergic action of the preferential detachment and of the geometrical hindrance on the *r*_*ATP*_(*ν*) relationship shown in this paper goes beyond the qualitative effects recognized explicitly or implicitly by previous works.

In conclusion, our theoretical and experimental results make a step toward the elucidation of non-conventional components in the chemo-mechanical transduction of energy in muscle, showing that not all ATP energy is available for the lever-arm swing.

## Materials and Methods

### Model

Conservation of the total number of myosin heads and of the actin target zones and their relative position is crucial for the analysis developed in this work. To overcome the intrinsic limits of mean-field models when accounting for these aspects, we use a Monte-Carlo approach. For each j-th thick filament, each i-th myosin motor position *x*_*i,j*_ and current state *s*_*i,j*_ are updated every time-step *Δt* depending on the conditions obtained from the previous time-step. Myosin motors are described by single material points in a coordinate system fixed with the relative thick filament and with the origin in the M-line.

We propose a pseudo-replication of the 3D structure of the sarcomere, neglecting the radial movements of the myosin motors and considering only the axial positions with respect to their anchor coordinates 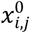, which replicate the geometrical hindrance as follows. Myosin motors belonging to the same crown (same color in Figure 1B and C) share the same axial anchoring position along the thick filament using the integer part of i/3, and crowns are spaced with a periodicity *d*_*myosin*_ = 14 3 nm. Allowing for ceil function symbology ⌈ ⌉, we have:

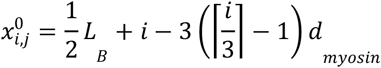

Along each half thick filament, a total of N_XB_=147 myosin dimers are distributed in this way [9]. In each crown, the three myosin dimers are rotated by 120°, and around each thick filament there are six thin filaments which run parallel to the thick filament, one every 60° (Figure 1B). Through our Monte-Carlo approach, we can use the actin filament periodicity to mimic the shift of the binding regions for every myosin motor due to the 3D structure (Figure 1B and C, same colors refer to binding regions for the corresponding motors). Actin filaments have a periodicity L_a_ of about 36 nm, and because of their double-stranded structure, the apparent periodicity is L_a_ every 180°, or L_a_/3 every 60° (Figure 1C). Imposing that the two heads in the first dimer of the first crown (*i* = 1) point precisely to one actin monomer of the corresponding thin filament (Figure 1B, deep red heads), then the other two dimers in the same crown point precisely to an(other) actin filament, but rotated relatively to the first thin filament by 120° each. This can be simulated with a target region (deep red regions on actin filaments in Figure 1B and C) shifted by 2/3 L_a_ and 4/3 L_a_ with respect to the first actin filament for the second and third dimers in the first crown, respectively.

Since we start with the first crown pointing directly to one actin filament, the second and third crowns will not point directly to their respective actin filaments (orange and yellow motors in Figure 1B and C) because of the mismatch between the dimer orientations (40° between consecutive crowns) and actin filaments (60° as noted above). We hypothesize that the heads can freely rotate in the azimuthal orientation (the polar angle with respect to the myosin backbone indicates the lever-arm rotation), so the ±20° will influence the attachment only through a shift of the target zones of ± L_a_/9, for the second and the third crowns, respectively, plus the above mentioned L_a_/3 shift for each 60° (colored regions on the actin filaments in Figure 1B). Thus, attachment is possible only if:

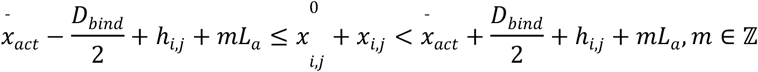

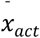 is the position of the actin filament that accounts for the relative sliding during isotonic contraction, *D*_*bind*_ is the length of the target zone, and *h*_*i*_ is the above mentioned shift of the target zone to mimic the 3D geometry, i.e. 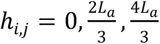 for i = 1 to 3 (first crown), 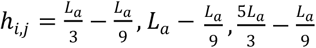 for i = 4 to 6 (second crown), and 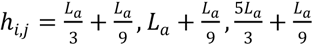 for i = 7 to 9 (third crown). Indices for crowns 4 to 49 are defined in the same way. When *D*_*bind*_ = *L*_*a*_, myosin motors can attach everywhere on the actin filament. In the text, this case is compared to 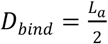 (three actin monomers are available to the attachment) and 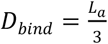 (only two actin monomers). The values of m are limited by the imposed length of the actin filament (1224 nm). To simulate variability in the initial orientations of the actin and/or myosin filaments, each target zone is shifted by L_a_ j/N_fil_, where j is the filament index and N_fil_ =120 is the total number of filaments considered.

Each myosin motor is subjected to thermal fluctuations and mechanical forces associated to a potential energy *E*(*x*_*i,j*_, *s*_*i,j*_) that depends on the myosin chemical state. Myosin motors can be in a detached super-relaxed, or OFF, state (*d*_*OFF*_), in a detached but active, or ON, state (*d*_*ON*_), and in an attached state in a pre-power stroke (*a*_0_), first power stroke (*a*_1_), or second power stroke (*a*_2_) configuration. Myosin motors are anchored to the thick filament through an elastic element of stiffness *K*_*m*_ = 2 pN/nm such that:

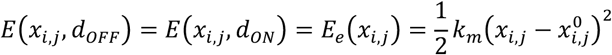

for both the ON and OFF detached states. The OFF state is probably more constrained than the ON state, but this point is not relevant in the current analysis, since the switch-ON rate does not depend on *x*_*i,j*_ (see below). Also, for some analyses specified in the text, the stiffness is asymmetric, with a value of 0.2 pN/nm when 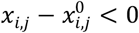.

When the actomyosin complex is formed, myosin motors are also subjected to a non-convex chemical potential energy with three minima corresponding to the three stable states (*a*_0_, *a*_1_, *a*_2_) defined as:

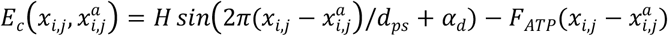

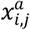 is the position of the myosin motor the moment it attaches to the actin filament, which defines the position of the minimum in the pre-power stroke state. H=5.7 *κ*_*B*_*T* (the Boltzmann constant times the absolute temperature) is the energy barrier between minima, *d*_*ps*_ =4.6 nm is the power stroke distance between stable state minima [24], and α_*d*_ is a constant value to impose the pre-power stroke minimum in 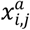. The last term is the bias induced by ATP toward the post power stroke states, with a decrease of 8 _*B*_ *T* every step in the power stroke. Thus, 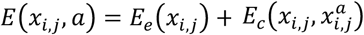 Quartic energy barriers are imposed outside the three minima.

In this work, we use the Kramers-Smulochovsky approximation of the corresponding Langevin equation driving the myosin dynamics, which is accurate for single-sarcomere and whole-fiber simulations, as previously shown [53]. In this framework, the effect of the thermal fluctuations is accounted by randomly selecting *x*_*i*_ from the stationary distribution of the probability density:

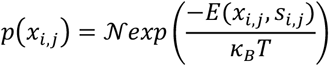

where *𝒩* is the normalization constant. The random selection is operated at any time-step by first mapping the probability such that 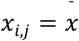 in a [0,1] range through the numerical integration:

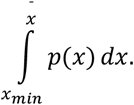

Then the position is defined at any time-step as the value that corresponds to a randomly generated number in the same range for the two detached states and for the three attached states.

The updated value of *x*_*i*_ is then used to update the value of *s*_*i*_ using a Metropolis algorithm, where a change from state “a” to state “b” with transition rate constant *K*_*a*-*b*_ occurs if a random number is in the range (0, *K*_*a*-*b*_Δ*t*), where *K*_*a*-*b*_Δ*t* ≪ 1 to ensure the generation of a Markov process.

Rate constants are as follows. OFF motors are activated with kinetics based on the experimental data obtained in [12]. OFF, or super-relaxed, motors switch to ON through the mechano-sensing mechanism described for the first time in [12], so 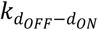 is a function of the force F_j_ sustained by the j-th thick filament. At any time-step, F_j_ is computed as:

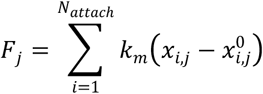

To mimic the experimental observations originally obtained by Linari and co-workers [12]:

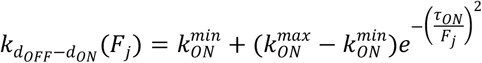

(see also [11]). The rate 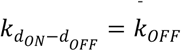 for active detached motors (ON) to switch OFF is supposed constant for simplicity.

ON motors can attach to an actin filament when the geometrical constraint allows in the *a*_0_ state, with a rate, which is a function of 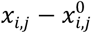, as proposed in the original Huxley model [6]:

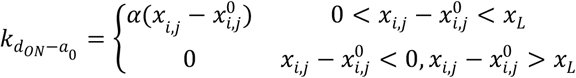

*x*_*L*_ = 10 nm as in the original model, even though in our Monte-Carlo approach 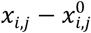 never reaches this value because of the elastic element constraint.

Detachment can occur in any attached state and is defined as:

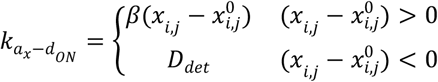

A mechanical detachment is imposed in the unlikely cases where the motor reaches the maximum limit of its S2 length when compressed (∼80 nm). Every detachment event leads to an ATP consumption, and this variable is also updated at any Δ*t*.

The probability for a motor to change state while attached is determined by the definition of 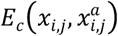 (in the Kramers-Smulochovsky approximation, which defines the rate constant from state *p* to state *q* as:

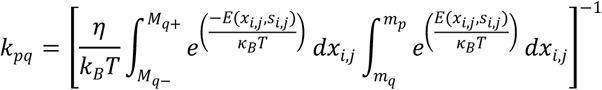

*M*_*q*+_ and *M*_*q*-_ are the right and left maxima, respectively, around the minimum energy for stable state *q* (*m*_*q*_), and the minimum energy for stable state *p* (*m*_*p*_). These values are computed at the beginning of the simulation and used at any time-step within the Metropolis algorithm.

The simulations are obtained for one half-sarcomere, and the fiber behavior is obtained under the hypothesis of a uniform contraction and force generation for each. The total force of the fiber is computed as 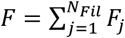, and 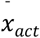 is updated following the isometric or isotonic external conditions with an implicit method (see [53]). The velocity is computed as

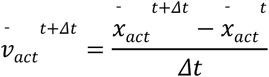

Finally, the total time is updated by summing Δ*t*, and the iterations continue until a given maximum time is reached.

### Construction of DNA origami

Ten helix-bundled DNA origami rods were designed using caDNAno software. To fold the DNA origami rod, 50 nM of scaffold (p8064, tilibit nanosystems) was mixed with 500 nM core staples. Oligonucleotides were obtained from Hokkaido System Science or IDT. The folding reaction was carried out in folding buffer (5 mM Tris pH 8.0, 1 mM EDTA and 18 mM MgCl_2_) with rapid heating to 80 °C and cooling in single degree steps to 60 °C over 2 h followed by additional cooling in single degree steps to 25 °C over another 72 h.

The folded DNA origami rods were purified by glycerol gradient ultracentrifugation. Briefly, 15–45% (v/v) gradient glycerol solutions in 1 × TE buffer containing 18 mM MgCl_2_ were made, and the glycerol fractions containing oligomeric DNA origami rods were determined by agarose gel electrophoresis. The concentration of the DNA origami rods was determined by a Nanodrop spectrophotometer (Thermo Scientific), and the solution was aliquoted and stored at −80 °C until use.

### Myosin construct

For wild type subfragment-1 (S1) of the myosin construct, human skeletal muscle myosin IIa cDNA (Kazusa Product ID FXC25901) was truncated at Ala849. This fragment included the motor domain, essential light chains (ELC) binding domain and regulatory light chains (RLC) binding domain. For oligonucleotide labeling and protein purification, SNAP-tag (New England Biolabs Inc.), FLAG-tag and 6 × His-tag were attached at the C-terminal via linkers (3 a.a., GGL). Two amino acids (Leu-Glu) corresponding to the restriction endonuclease recognition site (XhoI: CTCGAG) were kept between SNAP-tag and FLAG-tag. For the light chain null construct (lever-arm-less myosin S1), ELC and RLC binding sites (Lys786-Leu846) were deleted from the S1 construct. These myosin fragments were introduced downstream of the multi cloning site of the pShuttle-CMV vector (Agilent Technologies). For the double lever-arm length construct (double lever-arm length Myosin S1), the ELC and RLC binding domains were inserted just after Ala849 of WT S1.

### Protein expression and purification

Recombinant adenoviruses were produced using the AdEasy XL Adenoviral Vector System (Agilent Technologies). The produced adenoviruses were purified using the AdEasy Virus Purification Kit (Agilent Technologies). Recombinant myosin expression and purification were performed according to a previous study [24]. Briefly, murine C_2_C_12_ myoblasts (RIKEN Cell Bank) were cultured in DMEM (high glucose, Nacalai tesque) supplemented with 10% FBS (Gibco) and 1% Penicillin/Streptomycin (Nacalai tesque). To induce differentiation into myotubes, cells were grown to confluence, and the medium was replaced with DMEM supplemented with 2% horse serum (Gibco) and 1% Penicillin/Streptomycin. Forty-eight hours post differentiation, the cells were infected with 1 × 10^6–8^ plaque-forming units of virus. Forty-eight hours post infection, the medium was switched back to growth medium. After 3–5 days of medium exchange, the cells were washed with PBS and collected by cell scraping. The cells were then lysed with a dounce homogenizer and centrifuged. Recombinant myosin was purified from clarified lysate by using the AKTA purify system as follows (GE Healthcare). First, the myosin was purified by His-tag affinity purification with a 1 ml HisTrap HP nickel-sepharose column (GE Healthcare). The eluted myosin solution was then dialyzed overnight at 4 °C in a low-salt buffer (25 mM imidazole pH 7.0, 10 mM KCl, 4 mM MgCl_2_, 1 mM DTT). Finally, the recombinant myosin was purified on a 1 ml HiTrap Q HP sepharose anion-exchange column (GE) using a 0–1 M linear NaCl gradient.

### Oligonucleotide labeling to myosin

To label myosin to DNA origami, an oligonucleotide was attached to the SNAP-tag covalently just after anion-exchange purification. Amine-modified DNA oligonucleotides (NH2/GTGATGTAGGTGGTAGAGGAA) (Hokkaido System Science) were linked to the SNAP substrate, benzylguanine (BG; NEB), and 15–25 μM BG-oligonuculeotides were labeled with ∼1 μM myosin II containing a C-terminal SNAP-tag (NEB) in anion-exchange elution buffer for 30 min at room temperature. Oligonucleotide-labeled myosin II was purified by actin filament affinity to remove both unlabeled oligonucleotides and denatured myosins, aliquoted and stored at −80 °C until use.

### Labeling of Qdot to actin filament

Qdot 655 amine-derivatized polyethylene glycol (PEG) conjugates (4 µM; Invitrogen) were mixed with 50 mM Sulfo-SMCC (Thermo) and incubated for 1 h at room temperature. Excess Sulfo-SMCC was removed three times by gel filtration (Micro BIO- SPIN P-6, Biorad), mixed with thiol-modified DNA oligonucleotide (CTCTCCTCTCCACCATATCCA) and incubated overnight. Excess DNA was removed by Amicon-Ultra (100k NMWL, Merck), and the Qdot-oligonucleotides were stored at 4 °C until use.

The Qdot-oligonucleotides were mixed with a-actinin (actin binding protein) labeled with a complementary oligonucleotide at a molar ratio of 1:1 and incubated 2 h at 4 °C. ∼200 nM Qdot- actinin was mixed with ∼4 nM actin filament (filament concentration was calculated by assuming 2 µm in length) and incubated overnight. The Qdot-actin filament complexes were stored at 4 °C until use.

### Observation of actin sliding along DNA-origami thick filament

To avoid the nonspecific adsorption of myosins with the glass surface, a glass coverslip was coated with functional PEG according to our previous study [36]. Briefly, coverslips cleaned with a plasma cleaner for 10-15 min were soaked in freshly prepared 2% (3-Aminopropyl) trimethoxysilane (KBM-603, Shin-etsu Chemical) in acetone for 45 min with gentle shaking at room temperature. The coverslips were rinsed with ddH_2_O and dried. Next, 30 μl PEG solution (100 mg ml^−1^ (PEG 5000 (ME-0500HS, NOF Corp.):PEG 3400 (DE-034HS, NOF Corp.):biotin-PEG (BI-050TS, NOF Corp.)=100:10:1) in freshly prepared 0.1 M bicarbonate buffer pH 8.3) was put between a pair of coverslips and then incubated for 3 h. After rinsing with ddH_2_O, 20 μl of sulfodisuccinimidylartrate (Solteck Ventures) solution (30 mg ml^−1^ in freshly prepared 1 M bicarbonate buffer pH 8.3) was put between the pair of coverslips and incubated for 45 min. The coverslips were rinsed, dried and stored at −80 °C in a vacuum desiccator with desiccant.

A single flow chamber was made using double-sided transparent tape (Scotch) and biotinylated PEG-coated coverslips. Five microlitres of neutravidin (Invitrogen, 0.25 mg ml^-1^) was flowed into the chamber and incubated for 3 min. Unbound neutravidin was washed out by 2×Assay buffer (AB; 10 mM HEPES-KOH pH 7.8, 10 mM KCl, 4 mM MgCl_2_ and 1 mM EGTA). Blocking buffer (1 mg mg ml^-1^ BSA, 1 mg ml^-1^ casein in 50 mM KCl, 0.025% NaN_3_ and 10 mM Tris-HCl buffer pH 7.5) was flowed into the chamber and incubated for 4 min. Unbound BSA and casein were washed out by AB, and 0.1-0.5 nM biotinylated DNA-origami thick filament was flowed into the chamber and incubated for 3.5 min. Unbound thick filament was washed out by AB. 0.1-0.3 µM myosin was flowed into the chamber and incubated for 10 min. This step was repeated three times. Unbound myosin was washed out by motility buffer (MB; AB plus 0.5 µM ATP (Oriental Yeast), 0.436% ADP contaminants, an oxygen scavenger system and ATP regeneration system). ∼0.8 pM QDot-actin filaments in MB was flowed into the chamber, which was then sealed with nail polish and observed immediately. All experiments were performed at 24°C.

The observation of QDot-actin filament complexes was performed using TIRFM. Illumination was provided by 488 nm laser light (OBIS 488LS-100, Coherent) or 532 nm laser light (Compass 315M- 100, Coherent). The fluorescence of Qdot655 was passed through a dichroic mirror (FF552-Di02, Semrock) and a dual-view apparatus (Hamamatsu) equipped with dichroic mirrors (FF624-Di01, Semrock), which were put in front of the EMCCD camera (Andor, DV887ECS-BV). The fluorescent spots of QDot were fit to a 2D Gaussian function, and the center position was determined with 2-4 nm accuracy. The time trajectory was analyzed by a laboratory-written Labview program.

### Sliding filament model

The model for the DNA-origami is a modification of the fiber model to reproduce five repetitions of the basic element with 18 myosin heads spaced 14.3 nm apart. Contrary to the fiber model and following the experimental system geometry, the myosin motors are aligned along the same direction with no bare zone. There are no shifts due to different directions associated with the different crowns (*h*_*i,j*_ = 0∀*i, j*) such that:

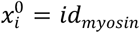

The OFF state, based on the myosin/backbone interaction, is not present in the experimental system and eliminated from the model. Other aspects and parameters are as defined for the fiber model and reported above. We simulated the sliding of 120 independent filaments and obtained the average velocity for three lever-arm lengths and for the zero lever-arm case. The latter is a simulation of no lever-arm rotation from the pre-power stroke state. The definition of the actomyosin energy 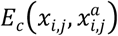 is the same as for the fiber model and is based on the step size *d*_*ps*_, therefore, we must define a correspondence between this parameter and the lever-arm length [54,55]. For the WT case, we impose two step-sizes of 4 nm each, as observed experimentally for the same

DNA structure from fast AFM images [24]. The no light chains case is simulated with two step sizes of 1 nm each [29] and the 4 light chains case with two step sizes of 7 nm each.

To reproduce the low [ATP] used in the sliding assay experiments, detachment of the myosin motor from the actin filament is modified to include an intermediate state *d*_*ON*-*ATP*_ that mimics the actomyosin complex after the release of ADP. From this state, the i-th myosin motor needs a new ATP molecule to detach and be available for a new cycle. 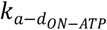 depends on the force acting on the actomyosin complex through the Bell equation [56]:

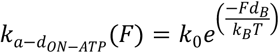

where *K*_0_=89 s^-1^ is the unloaded rate of detachment, and *d*_*B*_ =1.3 nm is the distance from the force dependent transition (data and values from [21]). 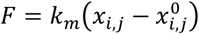 is the force.

The second step is then [ATP]-dependent, with a single constant rate 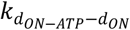 used to fit the sliding velocity in the WT.

## Acknowledgments

Funding was provided by the MSCA SoE @UniPD program (Acronym of the project: “Heart Fi-Re”, funding: Euros 150.000 to L.M.) from University of Padova, Grant-in-Aid for Young Scientists (B) (KAKENHI) (17K14375 to M.I.) from the Japan Society for the Promotion of Science (JSPS), and AMED-PRIME from the Japan Agency for Medical Research and Development (JP19gm5810022 to M.I.). We also acknowledge the CINECA award under the ISCRA initiative for providing high performance computing resources and support in the project IsC67_H-FiRe-1.

