## Supplemental text, figures and tables for "The synergic role of actomyosin architecture and biased detachment in muscle energetics: insights in cross bridge mechanism beyond the lever-arm swing"

**Supplementary Information for**
The synergic role of actomyosin architecture and biased kinetics in muscle energetics beyond the lever-arm swing.

Lorenzo Marcucci[*1,2], Hiroki Fukunaga[3], Toshio Yanagida[2,3,4], Mitsuhiro Iwaki[2,3]

1) Department of Biomedical Sciences, University of Padova, 35131 Padova, Italy
2) Center for Biosystems Dynamics Research, RIKEN, 5650874 Suita, Japan
3) Graduate School of Frontier Biosciences, Osaka University, 5650871 Suita, Japan
4) Center for Information and Neural Networks, NICT, 5650871 Suita, Japan

*corresponding author: Lorenzo Marcucci

**This PDF file includes:**

Supplementary text
Supplementary Figures S1 to S6
Supplementary Movie S1 caption
Tables S1 to S3
SI References

### Supplementary Information Text

**Single muscle fiber models:** The mathematical model used in the manuscript is based on a Monte-Carlo simulation with a time step $\Delta t$ of 1 μs. The parameters are reported in Table S1. The model is composed of $N_{fil}$ thick filaments, $L_M$ in length and each containing $N_{xb}$ myosin dimers arranged in crowns spaced by 14.3 nm starting after a bare zone $L_B$. Only one myosin motor can interact with the thin filaments at any $\Delta t$. Thin filaments are shifted as described in the Material and Methods in the main text to reproduce the 3D geometry and the emerging hinderance related to the limited actin target range. The model reproduces the behavior of a half-sarcomere and uses the hypothesis of homogeneous deformations to reproduce the experimental data from a single muscle fiber.

Thin filaments $L_A$ in length are regulated by the calcium concentration [$Ca^{2+}$] through Troponin-Tropomyosin units (TTU) in one of three states spaced every 36 nm. However, since the experimental data is based on fully activated muscle fibers, a very high [$Ca^{2+}$] is imposed for activation and kept constant during all the simulations. Full activation of the thin filament is reached in less than 1 ms. To avoid further parameters, the model doesn't include any cooperative effects on the myosin activation of the thin filament. These hypotheses are imposed to simplify the model and focus mainly on the dual filament activation through the mechano-sensing mechanism. Thermal fluctuations of the motors are computed both in the detached and attached states by considering a viscosity $\eta_{xb}$. A mechanical detachment is imposed in the rare cases where the motor reaches the maximum limit of its S2 length when compressed (-80 nm).

The benchmark models reported in Figure 3 without preferential detachment are not able to reproduce properly the tension-velocity curve or the kinetics of the tension generation and re-generation, mainly because $v_{max}$ is limited by the high viscous forces generated by attached motors in the compressed state. To partially solve this limit and obtain comparable results, we introduced a non-symmetric stiffness in the motors' elastic elements. Moreover, $r_{ATP}^{iso}$ is regulated by modulating the attachment and detachment rates and ignoring the rate of tension development. A detailed analysis of the influence of asymmetric stiffness in the actomyosin complex (1) on muscle energetics is beyond the scope of the present work (see for instance (2, 3)).

**Sliding filaments model**: The simulations for the sliding filament experiments are based on the same approach used for the muscle fiber but with the following modifications. Each thin filament acts independently and can slide in an unloaded condition driven by the force generated by the myosin motors. The OFF state is prevented and all motors are along the same line, so that the binding zones are defined from the same thin filament configuration. To improve the quantitative prediction of the model, an experimentally based detachment rate is imposed (see main text). This choice also imposes a modification to the attachment constants to better match the rate of force development/redevelopment and tension-velocity relationship in Figure S4 and the number of attached motors in isometric conditions. New parameters are reported in Table S3; the values $k_0$ and d refer to Bell's equation indicated in the main text.

Sliding filament experiments show slower kinetics compared to the fiber model. The simpler hypothesis for our simulations includes a dependence on the ATP concentration for the detachment of the myosin motor from the actin filament. After ADP release from the pocket, based on Bell's equation, 1 s is required to simulate the time needed for a new ATP molecule to reach the motor and permit detachment. This step is [ATP]-dependent through a single parameter, $k_{ATP}$, which in the sliding filament model is defined to fit the sliding velocity in the WT case.

The simulated sliding velocities fit well the experimental data, with a slightly lower match for the four light chains case (Figure 4C in main text). The longer lever-arm produces longer step sizes, and the required energy for a single step becomes higher than the drop in the energetic landscape (16 $\kappa_B T$ for the two steps (4)). In this situation, the slides should occur with clear steps, which is evidence of the coordinated motion of several attached motors (see SI figure S5). This behavior is not observed experimentally, probably because the longer lever-arm reduces its own stiffness (5). Assuming the drop in stiffness is linearly proportional to the length of the lever-arm, both the predicted stepwise motion (SI figure S5) and the simulated average velocity are more similar to what is observed experimentally (SI figure S6).

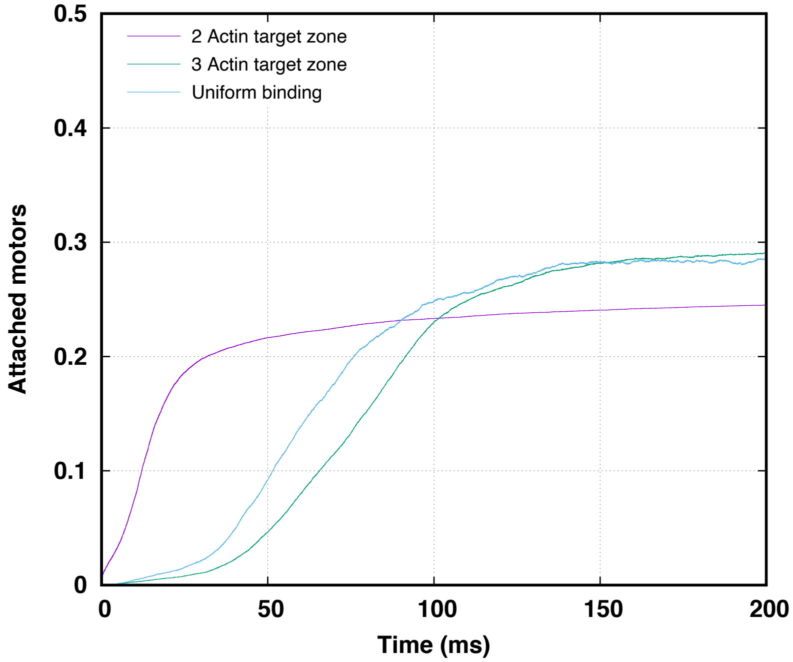


**Supplementary Figure S1**: Relative amount of strongly attached motors for the models with continuous binding sites (blue line), for limited binding regions of 3 actin monomers (green line) and 2 actin monomers 9purple line). In the first two cases, it is possible to fit the free parameters to match the experimental condition of 30% attached heads. In the case of only 2 actin monomers available for attachment, the maximum number of attached motors in isometric activation is limited to 25% by the geometrical hinderance. The maximum number of attached heads is obtained by fictitiously preventing detachment (shown by the monotonously increasing purple curve). Because of this unrealistic situation, the equilibrium is reached faster than the two other cases.

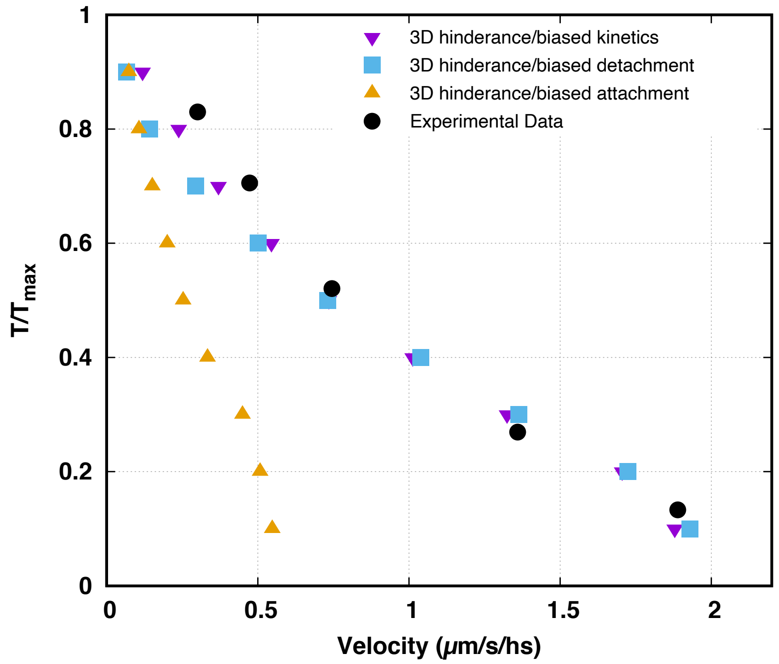


**Supplementary Figure S2:** Tension velocity relationships for the models with geometrical hinderance generated by limited target zones and preferential attachment and detachment (purple reversed triangles), preferential detachment only (blue squares) and preferential attachment only (orange triangles) and experimental data (black dots, see main text). Without preferential attachment, the tension-velocity curve can be sustained only with a faster attachment and detachment rate, leading to a higher ATP-ase consumption at intermediate velocities. It is possible to scale the maximum value of the ATP-ase rate by reducing the attachment rate, however, the simulated tension-velocity curve would show a slower velocity of contraction at intermediate tensions compared to experimental data. The preferential attachment keeps the tension-velocity curve inflection closer to the experimental data.

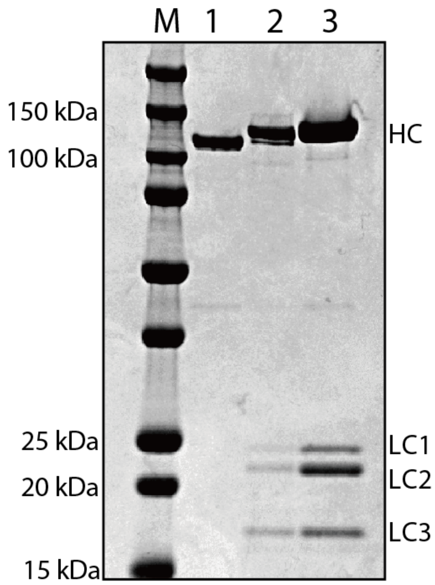


**Supplementary Figure S3**: SDS-PAGE gel of each myosin construct. Wild type (lane 2) and four light chains (lane 3) myosin express light chains (LC1-LC3, 15-25 kDa), while no light chain (lane 1) expresses no light chains. The bands around 100-150 kDa indicate heavy chains (HC). LC1 and LC3 are essential light chains, and LC2 is a regulatory light chain. Densitometric analysis of the bands revealed that the ratio of LC1+LC3 to HC was 0.8 and 1.8 for wild type and four light chains myosin, respectively, as expected from one or two essential light chain binding sites in the lever arm domain.
M, marker; lane 1, no light chain myosin; lane 2, wild type myosin; lane 3, four light chains myosin. HC, heavy chain; LC, light chain.

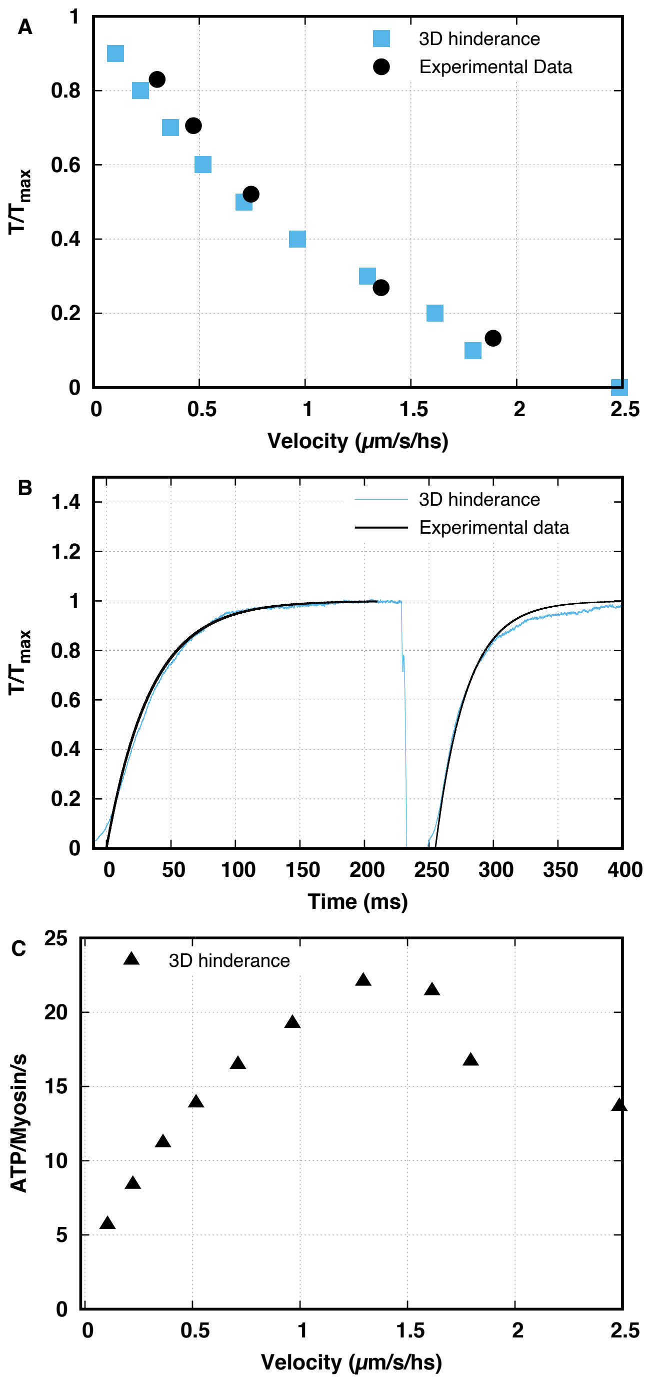


**Supplementary Figure S4**:
Fiber model simulations with the parameters used for the sliding filament model. A) Tension velocity relationships for the models with geometrical hinderance generated by limited target zones compared with experimental data (black dots, see main text). B) Kinetics of force development under isometric conditions and re-development after a small shortening at the unloaded velocity of contraction (blue line, simulations; black lines, experimental data from (6)). C) ATP consumption against the velocity of contraction.

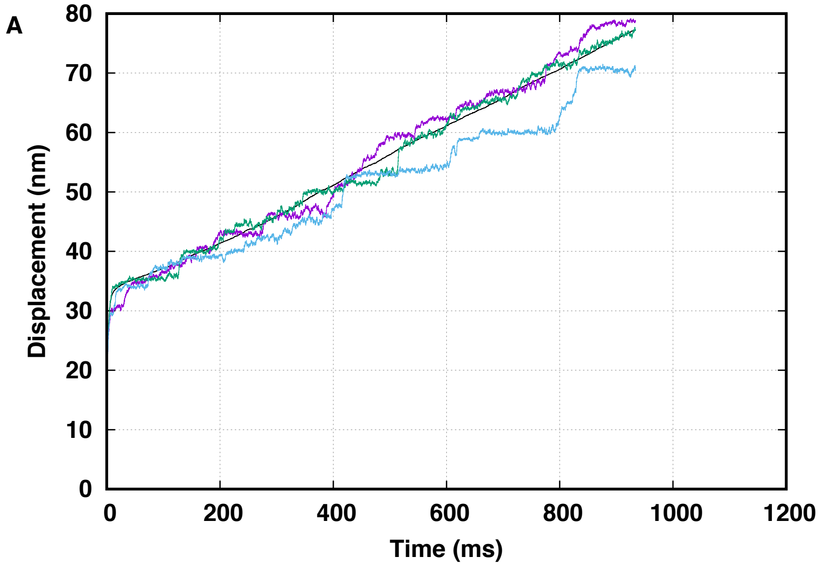


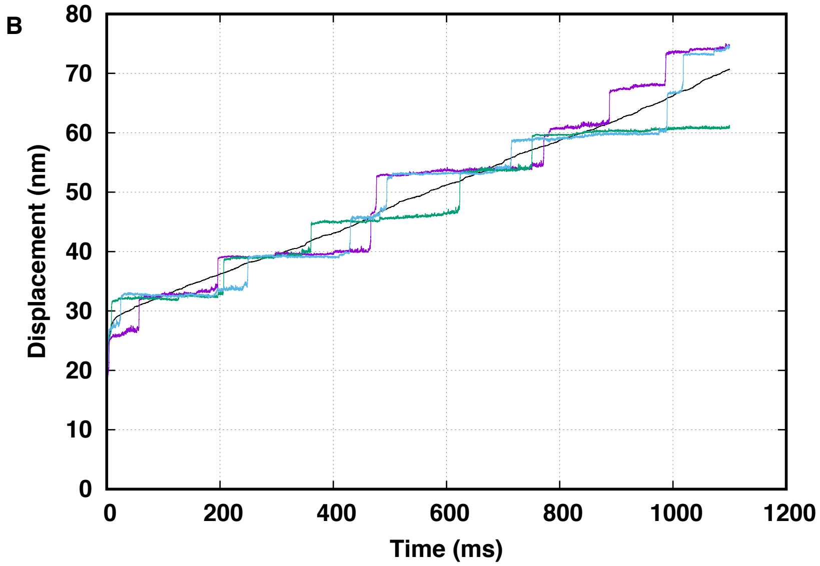


**Supplementary Figure S5:** Simulations of the mean trace (black) and typical single traces (colored) for the sliding of an actin filament in the DNA-origami model for the four light-chains case. A) An elastic element with a stiffness proportionally reduced to its length. B) An elastic element with a stiffness of 2 pN/nm.

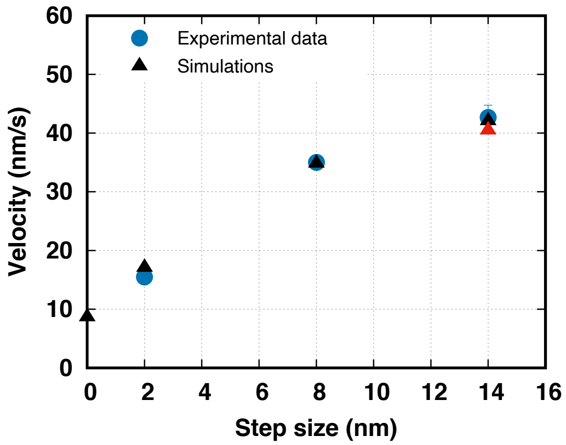


**Supplementary Figure S6:** Effect of the lever arm stiffness related to the lever arm length in the four light chains case. Predicted average velocities of the sliding model (triangles) at different step sizes and the corresponding velocities observed experimentally (blue dots) are related to different lengths of the lever-arm. The four light-chains model is shown at 2 pN/nm bending stiffness (red) and at a decreased stiffness due to the longer lever-arm (black).

**Supplementary Movie 1**: Actin sliding movement along a DNA rod. DNA rods (green) were sparsely adsorbed onto a glass slide, and a single actin filament (red) was unidirectionally moved along a DNA rod. ATP concentration, 1 μM. 10 Hz frame rate, x10 play.

**Table S1: Parameters for the muscle fiber models: common values**

| Parameter | Common value |
|---|---|
| $L_A$ | 1.2 μm |
| $L_M$ | 825 μm |
| $L_B$ | 50 μm |
| $N_{fil}$ | 120 |
| $N_{xb}$ | 147 |
| $\Delta t$ | 1 μs |
| $k_{xb}$ | 2 pN/nm |
| $\eta_{xb}$ | 70 pNns/nm |
| $\bar{k}_{OFF}$ | 196.2 $s^{-1}$ |

**Table S2: Parameters for the muscle fiber models: different values**

| Parameter | Uniform distribution | Geometrical hinderance (linear detachment model) | Geometrical hinderance (Bell's equation model) |
|---|---|---|---|
| $k_{ON}^{min}$ | 6.15 $s^{-1}$ | 2.8 $s^{-1}$ | 2.3 $s^{-1}$ |
| $\alpha$ | 129 $s^{-1}$ $nm^{-1}$ | 357 $s^{-1}$ $nm^{-1}$ | 852.8 $s^{-1}$ $nm^{-1}$ |
| $\beta$ | 10.4 $s^{-1}$ $nm^{-1}$ | 1.4 $s^{-1}$ $nm^{-1}$ | 1.4 $s^{-1}$ $nm^{-1}$ |
| $k_{ON}^{max}$ | 566.9 $s^{-1}$ | 566.9 $s^{-1}$ | 581.8 $s^{-1}$ |
| $\bar{k}_{OFF}$ | 196.2 $s^{-1}$ | 196.2 $s^{-1}$ | 342.3 $s^{-1}$ |
| $\tau_{ON}$ | 100 pN | 100 pN | 89.4 pN |
| $D_{det}$ | 610 $s^{-1}$ | 610 $s^{-1}$ | - |
| $k_0$ | - | - | 89 $s^{-1}$ |
| d | - | - | 1.3 nm |
| $k_{d_{ON-ATP}-d_{ON}}$ | - | - | 1400 $s^{-1}$ |

**Table S3: Parameters for the sliding filament model and the corresponding fiber model**

| Parameter | Sliding filament model |
|---|---|
| $L_A$ | 5 times 1.2 μm |
| $L_M$ | 5 times 825 μm |
| $L_B$ | 0 μm |
| $N_{fil}$ | 120 |
| $N_{xb}$ | 5 times 18 heads |
| $\Delta t$ | 1 μs |
| $k_{xb}$ | 2 pN/nm |
| $\eta_{xb}$ | 70 pNns/nm |
| $k_{ON}^{max}$ | 0 $s^{-1}$ |
| $k_{ON}^{min}$ | 0 $s^{-1}$ |
| $\tau_{ON}$ | 89.4 pN |
| $\bar{k}_{OFF}$ | 0 $s^{-1}$ |
| $\alpha$ | 852.8 $s^{-1}$ $nm^{-1}$ |
| $k_0$ | 89 $s^{-1}$ |
| d | 1.3 nm |
| $k_{d_{ON-ATP}-d_{ON}}$ | 4.2 $s^{-1}$ |
